# Drumming Performance and Underlying Muscle Activities in a Professional Rock Drummer with Lower-Limb Dystonia: A Case Study

**DOI:** 10.1101/2024.03.09.584247

**Authors:** Kazuaki Honda, Shizuka Sata, Mizuki Komine, Satoshi Yamaguchi, SungHyek Kim, Makio Kashino, Shinya Fujii

**Author notes:** **Correspondence:** Corresponding Author:, Shinya Fujii.

## Abstract

Task-specific focal dystonia (TSFD), characterized by the loss of fine motor control and coordination, affects drummers’ lower-limb movements, yet it remains relatively understudied, leaving a gap in understanding its effects on drumming performance and underlying muscle activities. This study explores lower limb dystonia’s impact on drumming performance and underlying muscle activity in a professional rock drummer. The drummer executed an eight-beat pattern on a drum kit, emphasizing kicking the bass drum on the initial beat and performing syncopation of the third beat. Dystonia symptoms primarily manifested in the initial beat, with fewer symptoms on syncopation of the third beat. Analysis revealed decreased bass-drum sound peak amplitude during the initial beat and increased (becoming more negative) synchronization error. Electromyographic (EMG) measurements of ten muscles in the affected right lower limb showed significant changes in the Biceps Femoris (BF), Tibialis Anterior (TA), Extensor Digitorum Longus (EDL), and Extensor Digitorum Brevis (EDB) muscles during symptom onset. Notably, we observed 1) earlier overactivation of the TA and EDL muscles during the leg lift-up motion or preparatory phase of pedaling, 2) reduced activation of the EDB muscle, and 3) increased activation of the BF muscle during the final pedaling movement when symptoms occurred. These findings suggest that lower-limb dystonia symptoms are characterized by a reduction in amplitude and an increase in synchronization error, potentially due to premature overactivation of the ankle dorsiflexor muscles.

## 1 Introduction

Task-specific focal dystonia (TSFD), characterized by the loss of fine motor control and coordination, significantly affects musicians’ careers (1). The body parts commonly affected include the right hand in pianists, the left hand in violinists (2), the embouchure in wind instrument players (3), and the lower limb in drummers (4–7). Reports on TSFD in drummers are limited compared with those on string, keyboard, and wind instrument players. There is a lack of knowledge regarding how dystonia affects drumming performance and the underlying muscle activities in drummers.

A previous study by Lee and Altenmüller (5) reported the drumming performance and underlying muscle activities in a patient performing a uniform accelerando motion with the lower limbs. In this study, the patient could perform this motion with the unaffected left leg up to a maximum speed of approximately 8 Hz, during which alternating muscle contractions were observed between the ankle plantar flexor and dorsiflexor muscles with minimal activation of the thigh muscles. Conversely, with the right leg affected, the patient could not maintain uniform acceleration, and the maximum speed for a stable rhythm was only 4–5 Hz. Notably, clear muscle co-contraction was observed in the thigh muscles of the affected right lower limb. Lee and Altenmüller (5) suggested that the control of drumming speed is compromised by lower-limb dystonia, and the co-activation of the thigh muscles is a characteristic of this symptom.

While the study by Lee and Altenmüller (5) provided valuable insights into the characteristics of lower limb dystonia symptoms in drummers, there have been no case reports on drumming performance and the underlying muscle activities when a drummer performs a pattern on a drum kit. Therefore, our study aimed to investigate these aspects in a professional rock drummer experiencing lower limb dystonia symptoms.

## 2 Methods

### 2.1 Participant

A 36-year-old male professional rock drummer participated in the study. He reported increased difficulty in his right lower extremity while playing with the drum kit. He began drumming at age 14 and practiced for 1–2 hours daily. After enrolling in a music college, he dropped out during his junior year and started his career as a professional drummer at the age of 20, practicing four hours a day. The first symptoms appeared at the age of 24 during a nationwide tour, manifesting as discomfort in the right foot. He experienced difficulty climbing stairs and driving when the symptoms were severe. To alleviate these issues, he adopted a ‘sensory trick,’ modifying his footwear and adjusting his chair height, which temporarily improved his symptoms. However, the relief was short-lived, and the effectiveness of the sensory trick diminished over time. He was diagnosed at age 29 by a neurologist. His family history was negative for any neurological disorders. Examination revealed a normal gait and no motor or sensory deficits in the lower limbs.

Ethical approval for this study was obtained from the Communication Science Laboratories Research Ethics Committee of Nippon Telegraph and Telephone Corporation (Approval Number: H30-009). The experiments were conducted in accordance with the principles of the Declaration of Helsinki. Written informed consent was obtained from the participant before the study.

### 2.2 Experimental setup and task

The experimental setup is illustrated in Figure 1a. The participant was instructed to play an eight-beat drumming pattern on a drumming kit at a tempo of 80 beats per minute (BPM), equivalent to 1.33 Hz, as illustrated in Figure 1b. The task was performed while listening to a pacing metronome-tone sequence via earphones. Each bar consists of four metronome tones, marked by the dashed vertical lines in Figure 1b, denoting the beats of the drumming pattern. The pattern involved playing a hi-hat, snare drum, or bass drum. Specifically, for the bass drum, the participant was required to strike two notes per bar, the first on the initial beat and the second on the syncopation of the third beat, represented by the orange and green notes in Figure 1b, respectively.

**Figure 1.**
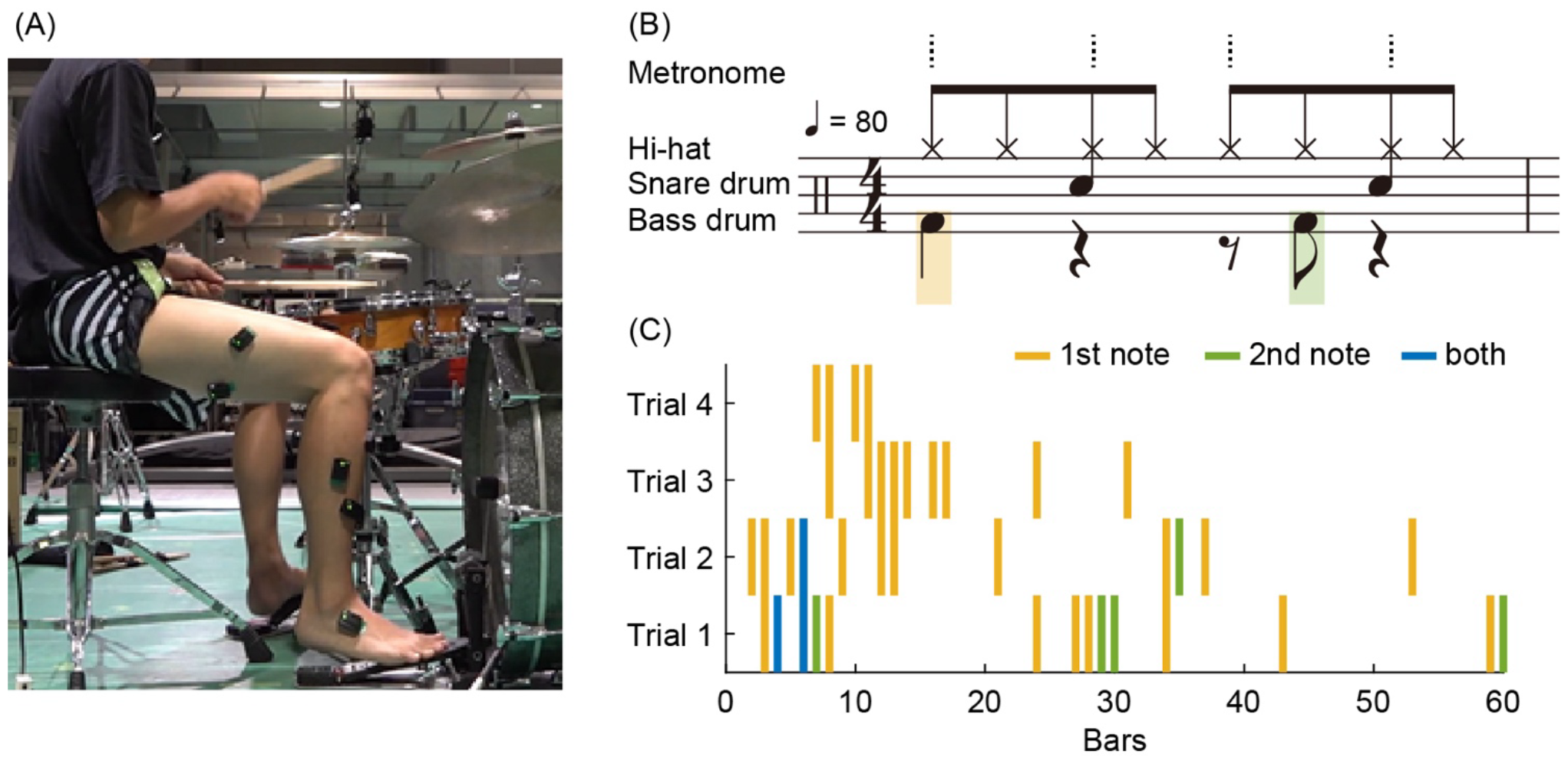
**(A)** Experimental setup: The participant was instructed to play an eight-beat drumming pattern on a drum kit at a tempo of 80 beats per minute (BPM) and to verbally report any occurrences of dystonia symptoms. **(B)** The pattern involved playing the high-hat, snare drum, and bass drum. For the bass drum, the participant was required to strike two notes per bar: the first on the initial beat and the second as a syncopation of the third beat, represented by orange and green notes, respectively. A bar consisted of four metronome tones, indicated by dashed vertical lines, denoting the beats of the drumming pattern. **(C)** Occurrence of dystonia symptoms across four trials: Forty-two instances of symptoms were reported; 31 occurred solely on the first note in a bar, 5 occurred only on the second note, and 3 occurred on both the first and second notes, as indicated by orange, green, and blue vertical lines, respectively.

The participant completed 60 bars in each trial. The start of a trial was signaled by a 1 kHz pure tone, followed by four metronome tones. The experiment comprised four trials. The participant was allowed a rest period of at least one minute between trials. During the trials, the participant was instructed to promptly and verbally report any occurrence of dystonia symptoms. We identified 42 instances of symptom occurrence (Figure 1c): 31 occurred only on the first note in a bar, five occurred only on the second note, and three occurred on both the first and second notes, indicated by orange, green, and blue vertical lines, respectively, in Figure 1c. This means that 34 instances (81%) occurred on the initial beat, and eight (19%) occurred on the syncopation of the third beat.

### 2.3 Data acquisition

A piezosensor (Model DT-10, YAMAHA) was attached to the head of the bass drum to record its impact. Simultaneously, we captured the piezosensor signal, metronome sound, and trigger signal for the Electromyographic (EMG) sensors using an audio interface (Fireface UCX, RME Audio) with a sampling rate of 48,000 Hz.

EMG activity was measured using active electrodes (Trigno IM sensor, Delsys Inc.). The EMGs recorded the muscle activities from ten muscles in the affected right lower limb: 1) Rectus Femoris (RF), 2) Vastus Lateralis (VL), 3) Vastus Medialis (VM), 4) Biceps Femoris (BF), 5) Tibialis Anterior (TA), 6) Extensor Digitorum Longus (EDL), 7) Soleus (SOL), 8) Gastrocnemius (GAS), 9) Peroneus Longus (PL), and 10) Extensor Digitorum Brevis (EDB). Each sensor was carefully placed on shaved skin to avoid innervation zones. The EMG signals were amplified, and the sampling rate was set to 1,111 Hz.

### 2.4 Data analysis

Owing to the recording failure of the EMG signals, the data from the first trial were excluded from the analysis. Consequently, the data from the last three trials, totaling 180 bars (60 bars × 3 trials), were analyzed. During these trials, 26 instances of symptom occurrence were recorded: 24 occurred only on the first note in a bar, one occurred only on the second note, and one occurred on both the first and second notes (as illustrated in Trials 2–4 in Figure 1c). Given that approximately 92% of the symptoms (24 out of 26 instances) occurred in the first note, our analysis primarily focused on the performance and EMG signals corresponding to the first beat in each bar. Specifically, of the initial 180 beats per bar in the last three trials, symptoms occurred in 24 beats, whereas no symptoms were observed in the remaining 156 beats.

To analyze the bass-drum and metronome sound signals, we identified the onset times as the points at which the envelope signals exceeded 10% of the peak amplitude of each burst. We calculated the synchronization error, defined as the time difference between the onsets of the metronome and bass drum, to assess the performance of the bass drum. In addition, the peak amplitude of the bass-drum sound recorded by the piezoelectric sensor was measured for each beat.

The EMG signals were synchronized with the piezo and metronome signals using the trigger signal. Initially, the EMG signals were rectified, followed by the calculation of the root mean square (RMS) using a 30-millisecond time window. To visualize muscle activation patterns with and without symptoms, we plotted the RMS of the EMG signals for a period of 750 ms before and after the onset of each metronome sound.

## 3 Results

### 3.1 Drumming performance

The peak amplitude of the bass-drum sound and the synchronization error during bass-drum play are illustrated in Figure 2. The light blue dots and lines represent data without symptoms (n = 156), whereas the orange dots and lines indicate data with symptoms (n = 26). The mean and standard deviation (SD) of the peak amplitude without symptoms were 0.30 ± 0.03 [arbitrary units (a.u.)], whereas with symptoms, they were 0.18 ± 0.09 [a.u.]. The mean amplitudes with symptoms were significantly lower than those without symptoms (Mann-Whitney *U* = 3404, *p* < 0.01, *r* = 0.45).

**Figure 2.**
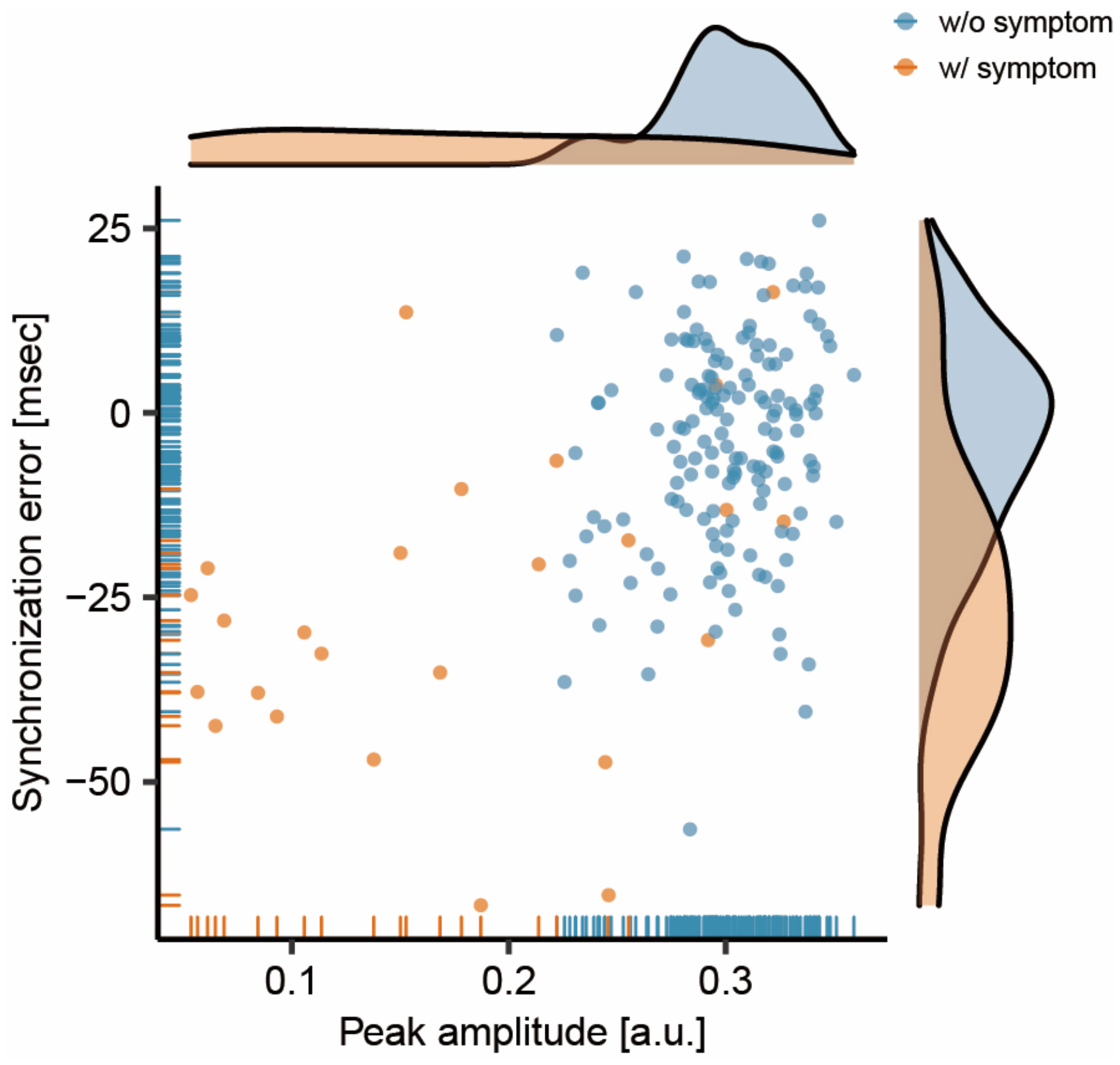
Relationship between the peak amplitude of the bass-drum sound and the synchronization error between the onsets of the metronome and the bass-drum sounds. The light blue dots and lines represent data collected without the presence of symptoms (w/o symptom), while the orange dots and lines indicate data collected with symptoms (w/ symptom).

The mean and SD of synchronization errors without symptoms were -4.02 ± 14.46 [milliseconds (msec)], and with symptoms, they were -26.22 ± 20.83 [msec]. The mean synchronization errors with symptoms were significantly larger (more negative) than those without symptoms (Mann-Whitney *U* = 3148, *p* < 0.01, *r* = 0.37). Overall, when symptoms occurred, the peak amplitude of the bass-drum sound decreased, and the synchronization error increased (became more negative).

### 3.2 Muscle activity

The RMS values of the EMG signals from the RF, VL, VM, BF, TA, EDL, SOL, GAS, PL, and EDB muscles are shown in Figure 3. The data were plotted for 750 milliseconds (msec) before and after the onset of each metronomic sound. The light-blue lines and areas represent the ensemble average and 95% confidence interval across the 156 instances without symptoms, whereas the orange lines and areas correspond to the 26 instances with symptoms. The heat maps display each of the RMS data sorted by the occurrence of symptoms and then by the degree of synchronization errors. Data above the red horizontal lines in the heat maps indicate instances of symptoms, whereas data below these lines represent those without symptoms. Data positioned higher on the heatmap correspond to EMG activity with larger synchronization errors (more negative errors).

**Figure 3.**
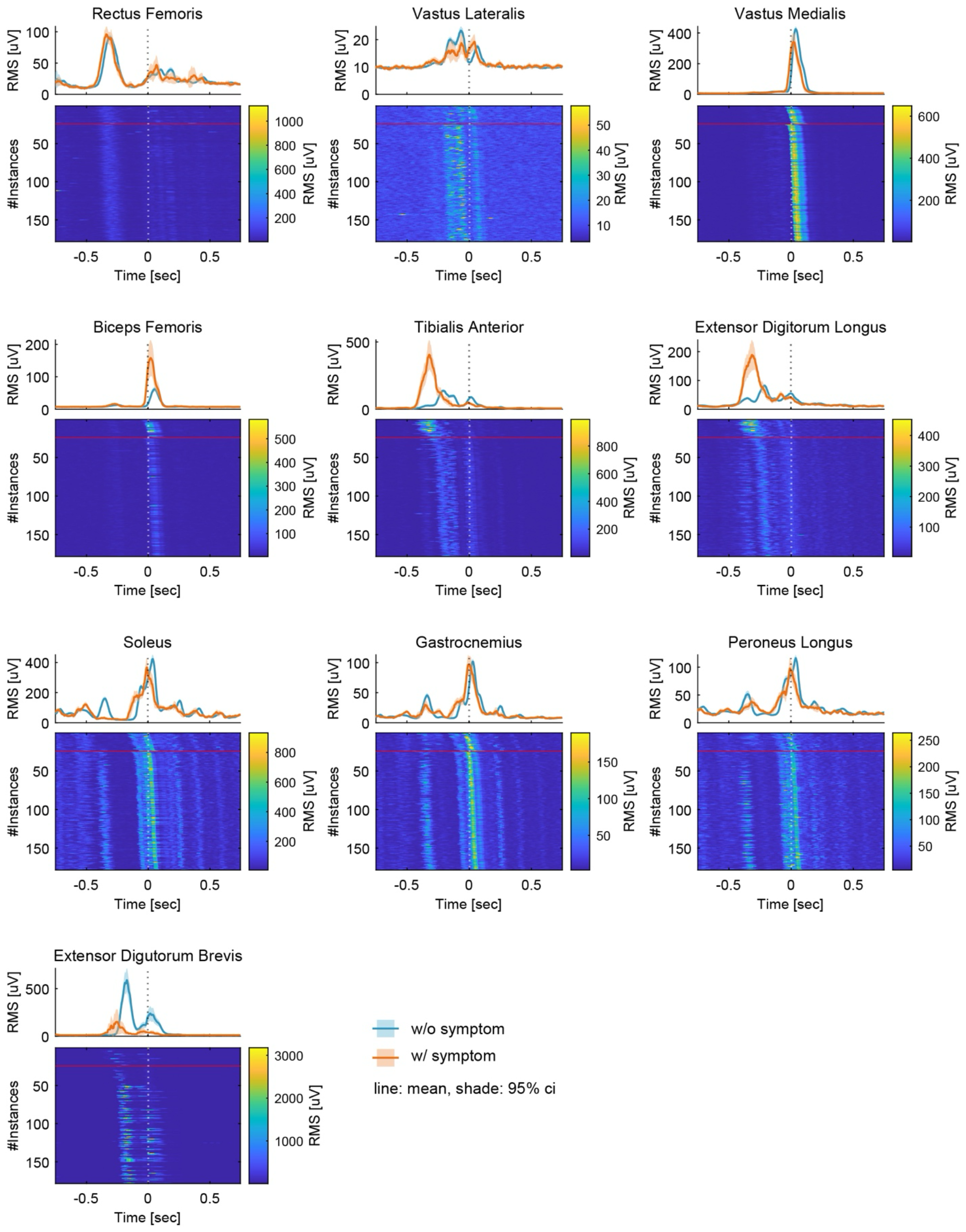
The root mean square (RMS) of the electromyographic (EMG) signals measured from 10 lower-limb muscles: Rectus Femoris (RF), Vastus Lateralis (VL), Vastus Medialis (VM), Biceps Femoris (BF), Tibialis Anterior (TA), Extensor Digitorum Longus (EDL), Soleus (SOL), Gastrocnemius (GAS), Peroneus Longus (PL), and Extensor Digitorum Brevis (EDB). Data were plotted for a period of 750 milliseconds (msec) before and after the onset of each metronome sound. Light blue lines and areas represent the ensemble average and the 95% confidence interval for instances without symptoms (w/o symptom), while orange lines and areas correspond to instances with symptoms (w/ symptom). The heatmaps display each set of RMS data, organized by the occurrence of symptoms and then by the degree of synchronization errors. In the heatmaps, data above the red horizontal lines indicate instances with symptoms, and data below these lines represent those without symptoms. Higher positions on the heatmap correspond to EMG activities with larger synchronization errors (more negative errors).

There were slight differences in the EMG patterns; however, the overall activities of the RF, VL, VM, SOL, GAS, and PL muscles were comparable between instances with and without symptoms. Conversely, notable differences were observed in the BF, TA, EDL, and EDB during the onset of symptoms. Specifically, TA and EDL activities shifted earlier and increased in magnitude, whereas the BF activity increased without a time shift. In contrast, the EDB activity shifted earlier and decreased when symptoms occurred.

## 4 Discussion

This study aimed to explore the impact of lower limb dystonia symptoms on drumming performance and the underlying muscle activities in a professional rock drummer. The drummer performed an eight-beat pattern on a drum kit, focusing on kicking the bass drum on the initial beat and syncoping with the third beat. The drummer reported the occurrence of dystonia symptoms, with most symptoms manifesting on the initial beat and a smaller percentage occurring on the syncopation of the third beat. Our analysis of drumming performance indicated that during the initial beat, there was a reduction in the peak amplitude of the bass-drum sound and an increase (becoming more negative) in the synchronization error. The EMG measurements of ten muscles in the affected right lower limb demonstrated notable differences in the Biceps Femoris (BF), Tibialis Anterior (TA), Extensor Digitorum Longus (EDL), and Extensor Digitorum Brevis (EDB) muscles during symptom occurrence. While a previous study (5) suggested that lower-limb dystonia compromises drumming speed control and is characterized by co-activation of thigh muscles, our findings indicate that the symptoms of lower limb dystonia are characterized by decreased amplitude and increased synchronization error, potentially due to earlier over-activation of the ankle dorsiflexor muscles.

### 4.1 Effect of dystonia symptoms on the downbeat

In our study, the drummer reported lower limb dystonia symptoms primarily on the initial beat, with fewer symptoms appearing upon syncopation of the third beat during an eight-beat drumming pattern. The symptoms were notably more pronounced on the downbeats (i.e., the count “one”) than on the upbeats (i.e., the syncopated beat following the count “three”). The predominance of symptoms during the initial beat may be attributed to factors related to attentional allocation and predictive timing mechanisms. According to the dynamic attending theory, attention is hierarchically allocated over time during the processing of hierarchical rhythmic structures (8–11). In the context of the eight-beat pattern used in our study, a strong attentional focus is likely directed towards the initial beat or the count “one.” Additionally, a neuroimaging study using magnetoencephalography (MEG) by Fujioka et al. (12) revealed that β-band event-related desynchronization is more pronounced at the downbeats compared to the upbeats. Fujioka et al. (12) suggested that the increased β-band activity at the downbeats is indicative of neural processing related to predictive timing or the conversion of timing information into auditory-motor coordination. Based on the findings of Fujioka et al. (12), we propose that the occurrence of dystonia symptoms in drummers may be linked to predictive timing mechanisms or the conversion of timing information into auditory-motor coordination.

### 4.2 Impact of dystonia symptoms on drumming performance

Our findings indicate a significant reduction in the peak amplitude of the bass-drum play when dystonic symptoms occur. The average peak amplitude with symptoms was measured at 0.18, compared to 0.30 without symptoms. This implied that the relative peak amplitude with symptoms was approximately 60% of that without symptoms, indicating a 40% loss in amplitude. Maintaining a strong bass-drum sound during the initial downbeat is essential for rock music performance. Previous research comparing skilled drummers with unskilled non-drummers found that skilled drummers demonstrated significantly less variability in tapping force, underscoring the importance of stable drumming performance (13,14). Therefore, it can be inferred that a reduction in peak amplitude may lead to a more variable performance, making it challenging for the drummer to maintain a steady rock beat.

Additionally, this study found that dystonia symptoms resulted in increased synchronization errors in bass-drum playing during the eight-beat pattern at 80 BPM. The average synchronization errors were -26.22 msec with symptoms and -4.02 msec without symptoms. A previous study investigating the synchronization error in drum kit playing at 60 and 120 BPMs by professional drummers reported average synchronization errors of -12.91 and -8.91 msec, respectively, across fifteen professional drummers (15). The synchronization error without symptoms in our study (-4.02 msec) was comparable to or even smaller than those reported in the previous study. However, with symptoms, the synchronization error (-26.22 msec) was significantly larger, approximately double or triple the average error observed in professional drummers in the previous study (-12.91 and -8.91 msecs). Therefore, the extent of synchronization errors with symptoms in our study was considered substantial, potentially impeding professional drummers’ ability to maintain precise timing.

### 4.3 Muscle activity in the presence of dystonia symptoms

In the absence of dystonia symptoms, sequential muscle activity associated with drum-pedaling movements was observed. Specifically, the rectus femoris (RF) muscle was activated for thigh lifting (hip flexion), whereas the gastrocnemius (GAS), soleus (SOL), and peroneus longus (PL) muscles extended the foot downward (ankle plantar flexion). The tibialis anterior (TA) and extensor digitorum longus (EDL) muscles facilitated upward foot extension (ankle dorsiflexion), while the extensor digitorum brevis (EDB) also contributed to toe extension. The vastus lateralis (VL) was active during knee extension, and the SOL, GAS, PL, vastus medialis (VM), biceps femoris (BF), and EDB were involved in the final striking action of the kick pedal.

In the presence of dystonia symptoms, three significant changes in muscle activity were observed: 1) earlier overactivation of the TA and EDL muscles during the leg lift-up motion or preparatory motion of pedaling; 2) subsequent diminished activation of the EDB muscle; and 3) overactivation of the BF muscle during the final pedaling movement. Premature overactivation of the TA and EDL muscles may cause excessive dorsiflexion of the ankle joint at an unintended earlier phase during symptom occurrence. Excessive dorsiflexion can impede subsequent toe extension, leading to reduced EDB muscle activation. Consequently, the drummer might activate the BF muscle and extend the hip to compensate for the movement, aiming to achieve the final striking action of the kick pedal. These unintended muscle activities and movements could result in an earlier and insufficient kick of the pedal, contributing to decreased amplitude and increased synchronization errors in drumming performance. These unintentional muscle activities, movements, and altered performance are likely the reasons why the participants could clearly report the occurrence of symptoms.

A previous study by Lee and Altenmüller (5) suggested that lower limb dystonia is characterized by the co-activation of the thigh muscles (BF and RF). In our study, we also observed relatively higher co-activation of the thigh muscles (BF, RF, VL, and VM) at the final kicking action, owing to the higher activity of the BF muscle when symptoms occur. However, the observed overactivation of the BF muscle may be a compensatory movement following the overactivation of the TA and EDL muscles. Thus, in our study, lower limb dystonia of the drummer might be characterized more by earlier overactivation of the ankle dorsiflexor muscles.

## 5 Acknowledgments

We thank T. Narita for her help with recording and editing the videos.

## 6 Data availability

The raw data supporting the conclusions of this article will be made available by the authors without undue reservation.

## 7 Author Contributions

All authors designed the experiments and collected the data. KH analyzed the data. KH and SF wrote the first draft of this manuscript. All authors reviewed and edited the manuscript and approved the final version.

